# Conjugation-mediated DNA delivery to the filamentous fungus *Ustilago maydis*

**DOI:** 10.1101/2025.11.13.688287

**Authors:** Tahani Jaafar, Amani Jaafar, Vida Nasrollahi, Bogumil J. Karas

## Abstract

Phytopathogenic fungi are ubiquitous throughout the environment and threaten global food security. This issue is further amplified by the increasing resistance of pathogens to antimicrobials. Current chemical-based antifungals target cells by inhibiting growth or metabolic function, making them ideal for fungal gain of resistance mutations. Biofungicides are a rising class of antifungals that have low potential for negative environmental impact and provide the fungi almost no potential for gaining resistance. Conjugative plasmids which play a role in the natural mechanism of horizontal gene transfer in bacteria, have been repurposed to deliver toxic genetic cargo to recipient cells, showing promise as next-generation antimicrobial agents. In this work, we have demonstrated the first protocol for delivering DNA from *Escherichia coli* to the filamentous phytopathogen, *Ustilago maydis* through conjugation. DNA delivery was confirmed using PCR screening of DNA isolated from the re-streaked transconjugants. Although challenges such as reduced conjugation efficiency and extrachromosomal replication persist, this work establishes the first step towards creating a conjugation-based biofungicide.

## INTRODUCTION

Pathogenic fungi are eukaryotic organisms that are widespread throughout the environment and cause significant medical and agricultural impact (Powers-Fletcher et al., 2016). Phytopathogenic fungi infect plants and cause disease, threatening global food security (Li et al., 2023; Möbius and Hertweck, 2009). This concern is amplified by climate change which has been suggested to be the cause of a global increase of their diversity and invasion potential. Researchers have estimated that the persistence of these diseases could leave enough food for only one-third of the world’s population, while plant diseases in forests may reduce global CO_2_ absorption annually by 230–580 megatons (Li et al., 2023).

Current methods of combating phytopathogens include the use of fungicides which are classified by their mode of action. Different modes of action include, damaging the fungal cell wall or cellular membrane, inactivating essential enzymes or proteins required for growth and stability, or interrupting critical processes for survival such as respiration and energy production (McGrath, 2004). Biofungicides are a novel class of treatments divided into three categories: 1) microbial biopesticides which contain a biological control agent that allows the organism to attack or compete with the phytopathogen, 2) plant biopesticides which are pesticides naturally produced from plant genetic material; 3) biochemical biopesticides which use natural material to disrupt the phytopathogen. Although the current treatments seem promising, the rise of fungicide resistance introduces a new challenge in the battle against pathogenic fungi. Fungicides with a single mode of action such as a large proportion of chemical fungicides are particularly at risk for the development of resistance. Multi-mode site of action fungicides have been developed but they have a higher potential in negatively impacting the environment including affecting non-target organisms (McGrath, 2004). Additionally, the decreased discovery of new fungicides and the increased demand have resulted in the perfect environment for fungal pathogens to gain resistance to the current treatments. This necessitates the development of novel fungicides that are at a lower risk of gaining resistance and have a low impact on affecting the environment.

*Ustilago maydis* is a filamentous basidiomycete that is a part of the Ustilaginaceae family, and causes smut disease in one of the world’s most substantial cereal crops, corn. The first high-quality assembly of the *U. maydis* genome was reported by (Kämper et al., 2006) with a genome size of 20.5 Mb that is made up of approximately 6900 genes. Later chromosome-level genomic assemblies reported 27 scaffolds interpreted as 23 chromosomes with a GC content of approximately 54% (Kämper et al., 2006). The dimorphic life cycle *U. maydis* begins with haploid sporidia (yeast-like growth) that develop into the pathogenic dikaryotic saprophytes (filamentous growth) after mating of compatible loci causing the fusion critical for infection and formation of the tumour-like galls on maize (Matei and Doehlemann, 2016).

Early genetic work identified an Autonomously Replicating Sequence (ARS) from *U. maydis* referred to as UARS, which enabled extrachromosomal replication of plasmids transformed into *Ustilago* (Tsukuda et al., 1988). However, the stability of the plasmid was shown to be limited due to the lack of a centromere (CEN) element that would provide mitotic stability. Genetic tools developed for *U. maydis* include integrative transformation systems, a range of selection markers (hygromycin, geneticin/G418, nourseothricin, carboxin), expression cassettes driven by native promoters (e.g. GADPH promoter), and advanced genome editing via CRISPR/Cas9 systems (Li et al., 2017; Schuster et al., 2016). In addition, various transformation methods were developed including protoplast-mediated (PEG) transformation (Wang et al., 1988), electroporation (Fischer, 2002), and *Agrobacterium tumefaciens*-mediated transformation (ATMT) (Ji et al., 2010).

Largely driven by horizontal gene transfer (HGT), life on earth is resultant of a vast, interconnected network of genetic exchange, rather than isolated lineages. Among the mechanisms that drive this exchange is bacterial conjugation (Tatum and Lederberg, 1947). Further research elucidated conjugation as an evolutionarily conserved, contact-dependent method of DNA delivery among gram-positive and gram-negative bacteria. Most conjugative systems utilize a Type IV secretion system (T4SS), with the structure and composition of the T4SS macromolecules varying between different plasmid groups (Christie, 2001). DNA delivery is dependent on presence of a conjugative plasmid encoding the necessary machinery. HGT, particularly through conjugation, drives the rapid spread of genetic traits like antibiotic resistance, virulence, and metabolic capacity across microbial communities. Conjugation was initially believed to be limited to prokaryotes, but it has been demonstrated that conjugation can deliver DNA into eukaryotic species like plants, algae, and yeast highlighting the potentiality for inter-kingdom DNA transfer and suggesting broader host ranges than previously realized (Bates et al., 1998; Brumwell et al., 2019; Cochrane et al., 2022; Hayman and Bolen, 1993; Karas et al., 2015; Moriguchi et al., 2013; Soltysiak et al., 2019; Suzuki et al., 2015). Harnessing bacterial conjugation for fungal DNA delivery presents a promising approach to overcoming host-specific barriers, expanding the opportunities for genetic manipulation in fungi as well as development of antifungals (Cochrane et al., 2022).

Our newly developed conjugation-based delivery method provides an alternative route for DNA transfer into *U. maydis*, potentially overcoming limitations of other delivery methods by enabling direct inter-kingdom transfer of plasmids without protoplast formation or strain-specific optimization. This platform establishes a foundation for future applications, including conjugation-driven antifungal systems that could terminate fungal cells by introducing toxic genes or deleting essential loci via CRISPR (similarly as done for *S. cerevisiae*, Nasrollahi et al, manuscript in preparation). Furthermore, conjugative plasmids can incorporate inducible control systems to enhance biocontainment and regulate DNA transfer (Jaafar and Carvalhais, 2025). In this paper we have demonstrated the first step towards this goal, successful conjugation from *E. coli* to the filamentous phytopathogen, *U. maydis*.

## MATERIALS AND METHODS

### Strains and growth conditions

*Escherichia coli* Epi300 (Lucigen Corp., Cat #: LGN-EC300110, USA) carrying the conjugative plasmid was grown in Luria-Bertani (LB) media supplemented with gentamicin (40 µg/mL; BioBasic, Cat #: GB0217, Canada. Recipient *Ustilago maydis* (Canadian Collection of Fungal Cultures, DAOMC 251941M) was grown in 2x yeast extract peptone dextrose media supplemented with adenine hemisulfate (200 µg/mL; Sigma–Aldrich, #A2545, St. Louis, MO, USA) (YPAD) and ampicillin (100 µg/mL; BioBasic, Cat #: AB0028, Canada).

### Conjugative plasmids

pSC5GGv1 plasmid was created by (Cochrane et al., 2022). pSC5-UARS was created using GoldenGate by inserting BsaI cut sites while amplifying the UARS region from the pGL0_21 [UARS1 ORI] plasmid gifted from Charlie Gilbert & Nili Ostrov & Cultivarium Tools (Addgene plasmid # 198934; http://n2t.net/addgene:198934; RRID:Addgene_198934) (Gilbert et al., 2023). Primers and templates used to amplify all the fragments are listed in Supplemental Table S1. pTAMob2.0 plasmid was created by (Soltysiak et al., 2019).

### Conjugation to *U. maydis*

#### Preparation of E. coli

Donor *E. coli* was grown overnight at 37 °C with shaking at 225 rotations per minute (rpm) in 5 mL of LB media inoculated with a single colony and supplemented with gentamicin. Three *E. coli* strains were grown containing one of three plasmids: pSC5GGv1, pSC5-UARS, or pTAMob2.0. Saturated cultures were diluted to an optical density at 600 nm (OD_600_) of 0.1 into a 50 mL culture of LB the following day and supplemented with gentamicin. The 50 mL cultures were grown to an OD_600_ of 1.0 and pelleted for 15 min at 3000 Relative Centrifugal Force (RCF). After centrifugation, the supernatant was decanted, and the pellet was resuspended in 500 uL of ice-cold 10% glycerol. Aliquots of 200 µL were frozen in a −80 °C ethanol bath and stored at −80 °C.

#### Preparation of U. maydis

Recipient *U. maydis* was grown overnight at 30 °C with shaking at 225 rpm in 5 mL of 2x YPAD media inoculated with a single colony and supplemented with the ampicillin. Saturated cultures were diluted to an OD_600_ of 0.1 into a 50 mL culture of 2x YPAD the following day and supplemented with ampicillin. The 50 mL cultures were grown to an OD_600_ of 1.0 and pelleted for 15 min at 3000 RCF. After centrifugation, the supernatant was decanted, and the pellet was resuspended in 1 mL of ice-cold 10% glycerol. Aliquots of 100 µL were frozen in a −80 °C ethanol bath and stored at −80 °C.

#### Conjugation

Conjugation plates (25 mL, 1% agar, 2x YPAD, no antibiotic) were prepared. Aliquots of the donor and recipient were thawed on ice for approximately 20 minutes. Then, 200 µL of the donor cells were added to 100 uL of the recipient cells in an Eppendorf tube and pipetted up and down before adding the total volume to the conjugation plate and spreading the mixture evenly. Plates were allowed to dry for 5 min before incubating at 30 °C for 3 h without stacking. Following the conjugation period, plates were scraped with 1 mL of cold sterile ddH_2_O and readjusted in an Eppendorf tube to a total volume of 1 mL. Next, the cell suspension was vortexed for 30-60 seconds, or until homogenous. The undiluted cell suspension was then spread on selection plates (30 mL, 1% agar, 2x YPAD with ampicillin (100 μg/mL) and nourseothricin (50 or 100 μg/mL; Jena BioScience, Cat #: AB-102XL, Germany)). Plates were allowed to dry for 5 min before being incubated at 30 °C for 6-11 days.

### DNA isolation from *U. maydis* transconjugants

DNA were isolated from all three plasmid transconjugants in both colonies presented in Figure 3 using a modified Cetyltrimethylammonium bromide (CTAB) protocol as previously described (Walker and Karas, 2025). First, the transconjugant was re-streaked on to a 2x YPAD plate supplemented with ampicillin and nourseothricin (50 μg/mL) and grown for 7 days, 3 times to ensure no *E*. coli were remaining. Next, the final re-streaked patch of approximately 1cm x 1cm was picked from the plate and dropped into liquid nitrogen until frozen. Next, the NEB Monarch High Molecular Weight DNA Extraction kit (NEB #T3060) was used by inserting the frozen re-streak into a Monarch Pestle Tube (NEB #T3001) and ground with the Monarch Pestles (NEB #T3002) until a fine powder was achieved. In the case of the re-streak thawing too quickly, the tube was closed and dropped in liquid nitrogen to re-freeze before resuming the grinding process. The tubes were then spun down to pellet the powder, 500 uL of CTAB lysis buffer, 0.5 uL of RNase A (QIAGEN, catalog number: 19101), and 1 μL (i.e., 100 μg) of proteinase K solution (20 mg/mL, BioShop, catalog number: PRK222) were added. The tubes were then slowly mixed using end-over-end inversion, and incubated at 37 °C for 15 min. The tubes were then gently mixed using end-over-end inversion once more, and placed into a 37 °C incubator for another 15 min. Next, the cell resuspension was pelleted by spinning at 16,000 RCF for 5 min in a microcentrifuge. The lysate was transferred to a new 1.5 mL Eppendorf tube, and one volume of phenol:chloroform:isoamyl alcohol (25:24:1) was added and mixed gently by end-over-end inversion. Next, the sample was centrifuged at 16,000 RCF for 5 min, and the aqueous phase (i.e., top layer) was transferred to a new 1.5 mL Eppendorf tube. One volume of chloroform:isoamyl alcohol (24:1) was added and mixed gently by end-over-end inversion. The sample was then centrifuged at 16,000 RCF for 5 min. Next, approximately 450 μL of the aqueous phase was added to a new 1.5 mL Eppendorf tube. To the this tube, 45 uL of sodium acetate solution (3 M, pH 5.2) and 9 uL of ice-cold 100% ethanol were added and mixed gently using end-over-end inversion. The sample(s) were then left overnight in the −20ºC freezer to increase the yield of DNA. The following day, the sample(s) were centrifuged at 16,000 RCF for 5 min, and the supernatant was decanted. The tubes were inverted on a paper towel to dry until all residual ethanol had evaporated. Finally, the pellet was resuspended with 30 μL of pre-warmed sterile ddH_2_O and store at −20 °C. The concentration and purity of the genomic DNA was measured using a spectrophotometer (Supplementary Table S2).

### Single-plex PCR screening for DNA delivery

Single-plex PCR screening was performed with GXL polymerase (Takara, catalog number: R050A) using the rapid protocol and Primer3-optimized primer pairs (6 primers in total; Supplementary Table S3). 25 µl reactions were performed using 1 µl of genomic DNA isolated from the transconjugant and diluted to 10 ng/uL. For all screens, the positive control consisted of ∼10 ng of purified pSC5GGv1 plasmid DNA and the negative control consisted of 1 µl ddH_2_O.

### Multiplex PCR screening for DNA delivery

Multiplex PCR screening was performed with Qiagen Multiplex Kit (Qiagen, Inc., Cat #: 206143, Germany) according to the Qiagen Multiplex PCR Handbook and Primer3-optimized primer pairs (6 primers in total; Supplementary Table S4). 10 µl MPX reactions were performed using 1 µl of genomic DNA isolated from the transconjugant and diluted to 10 ng/uL. For all screens, the positive control consisted of ∼10 ng of purified pSC5GGv1 plasmid DNA and the negative control consisted of 1 µl ddH_2_O.

### Statistical analysis

A two-way repeated-measures ANOVA test was ran (within-replicate factors: **plasmid** with 3 levels, and **concentration** with 2 levels). Next, a paired t-test was performed for all pairwise comparisons (comparing the three plasmids at each concentration of antibiotic, and a low vs. high concentration of antibiotic for each plasmid). This resulted in nine tests. Data were expressed as mean ± standard error of the mean of eight biological replicates. Bonferroni’s correction was applied to these paired tests, where the tests were considered statistically significant when P < 0.005 (^*^). All statistical values displayed in Supplementary Table S5, S6.

## RESULTS

To test conjugation as a method of DNA delivery to *U. maydis*, we used pSC5GGv1 (56.9 kb), which contains all necessary conjugation machinery and has been shown to be delivered to diverse fungi. Additionally, it contains the yeast backbone compromised of an ARS for *S. cerevisae*, a CEN element, and an auxotrophic selective marker (Cochrane et al., 2022). In addition we created the plasmid pSC5-UARS (56.1 kb) which contains the ARS isolated from *Ustilago, UARS* (Tsukuda et al., 1988). As a negative control, we used the plasmid pTAMob2.0 (Soltysiak et al., 2019) which contains an origin of transfer (oriT) as do the two previously described plasmids but lacks the nourseothricin resistance cassette which both pSC5GGv1 and pSC5-UARS contain. Figure 1 displays plasmids maps and the experimental design.

**Figure 1.**
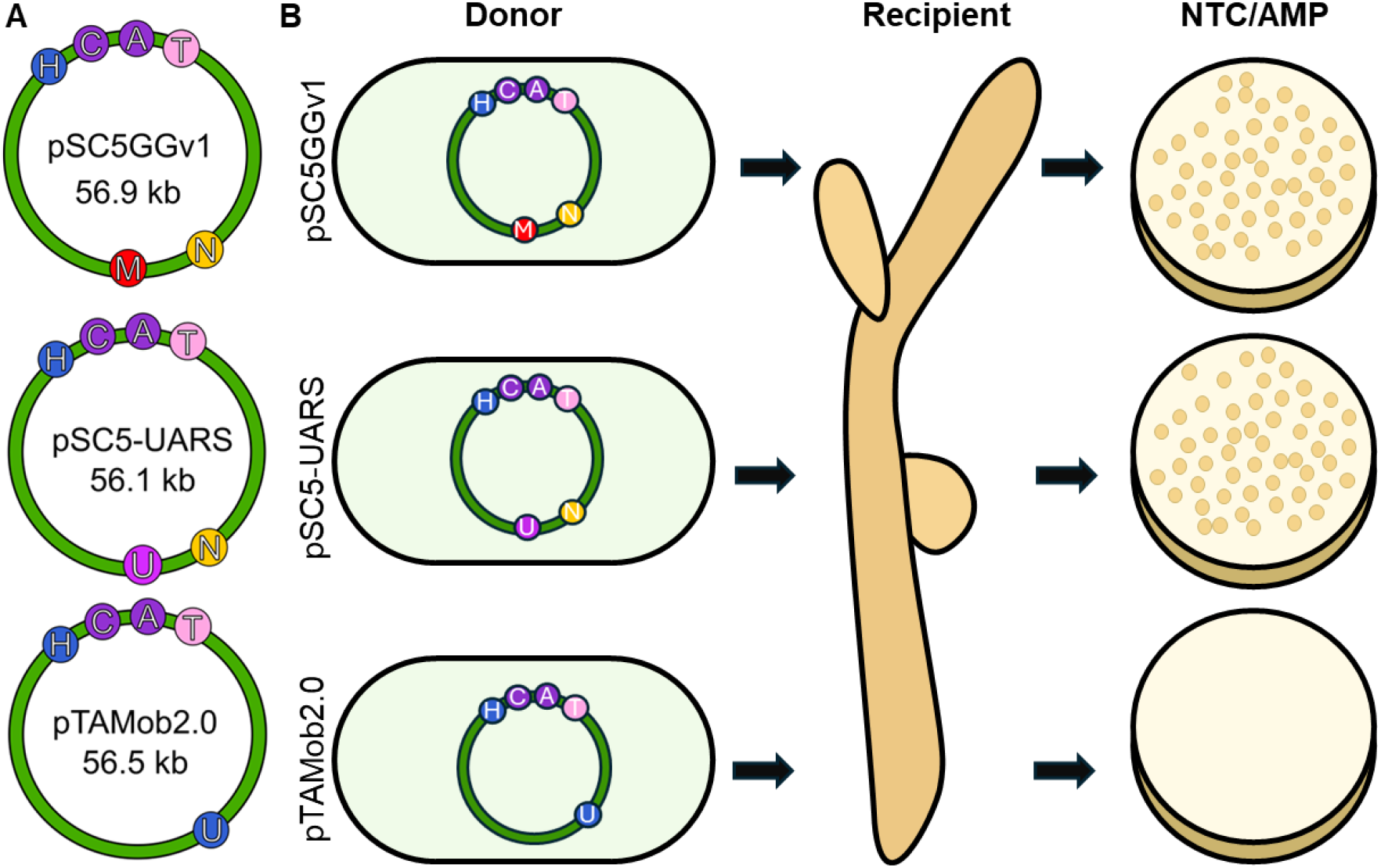
Schematic of plasmids used in *cis*-conjugation to *U. maydis* and possible outcomes. (A) All plasmids contain the auxotrophic yeast selection marker displayed in blue, HIS3 and pTAMob2.0 (Soltysiak et al., 2019) additionally contains URA3. The plasmid maintenance elements, CEN and ARS4 are displayed in purple, and the oriT in pink. pSC5GGv1 contains a mRFP gene displayed in red. pSC5-UARS contains the *UARS* sequence, displayed in magenta in place of the mRFP. Both pSC5GGv1 and pSC5-UARS contain the nourseothricin resistance cassette. All three of these plasmids have gentamicin resistance as a selectable marker in bacteria. (B) Schematic of possible outcomes following the *E. coli* to *U. maydis* conjugation. Using selection plates containing nourseothricin and ampicillin, it is possible to evaluate the transfer of the conjugative plasmid.

We based our initial conjugation protocol on the protocol published by (Cochrane et al., 2022). However, our initial results were not consistent, therefore we performed additional experiments to optimize the protocol for *U. maydis* by testing different ratios of bacteria to fungi (Supplementary Table S7) and different types of media for the conjugation plates (Supplementary Table S8). This yielded the final protocol described in the Materials and Methods section that was reproduced in eight biological replicates (Figure 2). The earliest appearance of colonies was approximately between 4-6 days of incubation where all colonies appear to be of similar size. We tested colony growth at two different nourseothricin concentrations (NTC 50 µg/mL and NTC 100 µg/mL) and found that by decreasing the antibiotic concentration, we achieved a larger number of colonies but with visible breakthrough using the control pTAMob2.0 plasmid (Figure 2A). However, increasing the incubation period unveiled two distinct colony sizes which we describe as ‘small’ and ‘big (Figure 2B). The average of the biological replicates across the different antibiotic concentrations showcased no significant difference between using pSC5GGv1 as compared to pSC5-UARS (Figure 2C). However, both plasmids had significantly more colonies when compared to pTAMob2.0, at both tested concentrations of nourseothricin. Overall, we identified two conjugative plasmids (pSC5GGv1 and pSC5-UARS) which resulted in colony growth, and when plated on the lower concentration of nourseothricin two clear colony morphologies can be identified. The small colony is approximated to be 10-20 times smaller than the big colony, as well as having a harder, smoother appearance with white colouration. The big colony exhibited a larger size with a softer, fuzzy appearance and darker yellow colouration.

**Figure 2.**
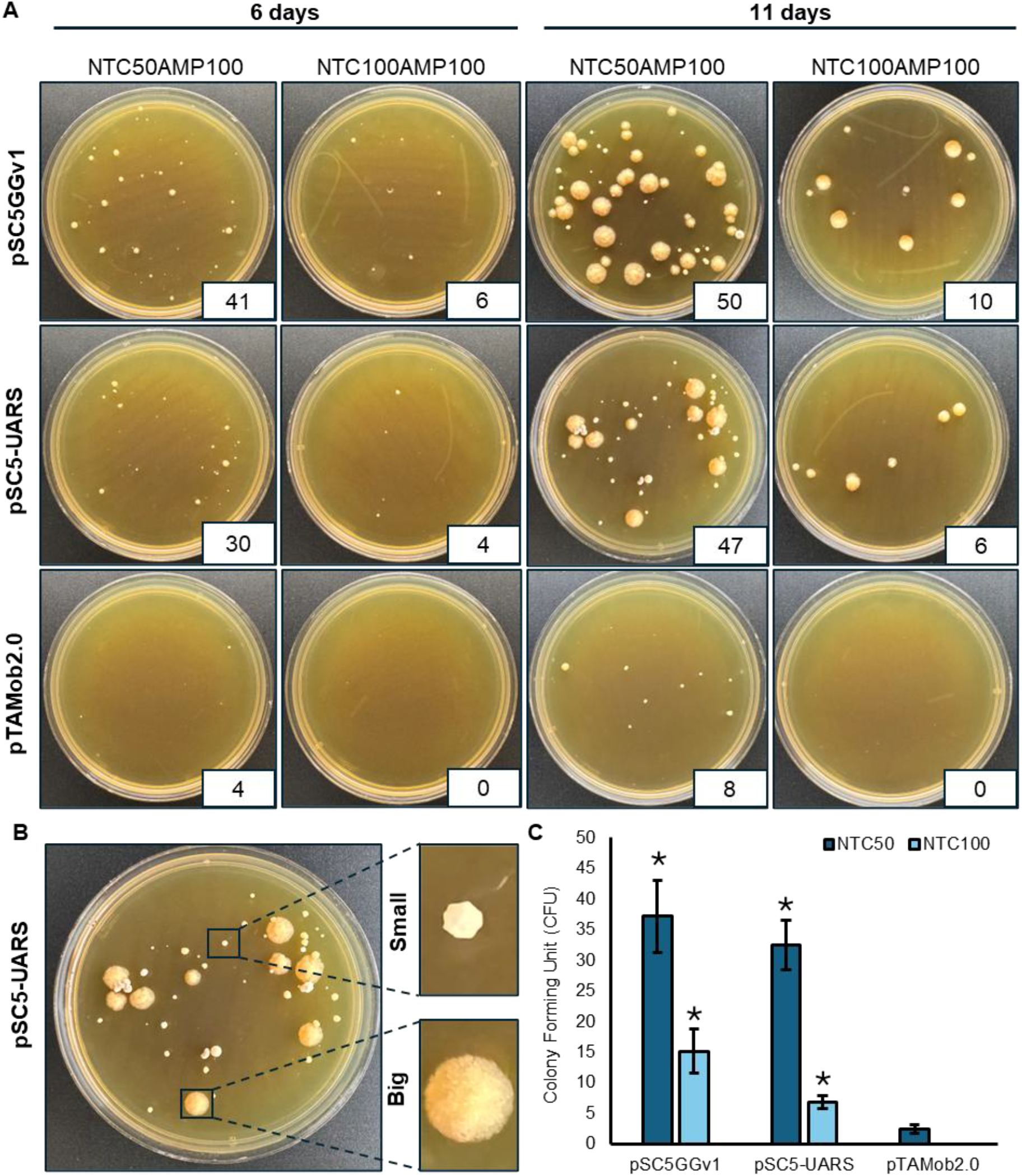
Analysis of the conjugation to *U. maydis*. (A) Representative transconjugant colonies on final selection plates (NTC/AMP). The three donor strains are displayed on the left: pSC5GGv1, pSC5-UARS, and pTAMob2.0. Selection on two different antibiotic concentrations: NTC 50 µg/mL/AMP 100 µg/mL and NTC 100 µg/mL/AMP 100 µg/mL. Complete figures of all replicates displayed in Supplemental Figure S1. (B) Representative plate from the pSC5-UARS conjugation selected on NTC 50 µg/mL/AMP 100 µg/mL and grown for 11 days. Two distinct colony morphologies are displayed: ‘small’ and ‘big’. (C) Average colony counts across all eight biological replicates for plasmid delivery at both nourseothricin concentrations. Error bars represent the standard error of the mean (SEM). Paired t-test with Bonferroni’s correction: ^*^**p** < 0.005, n= 8. The asterisk represents a significant difference between the respective plasmid and pTAMob2.0 at each concentration.

**Figure 3.**
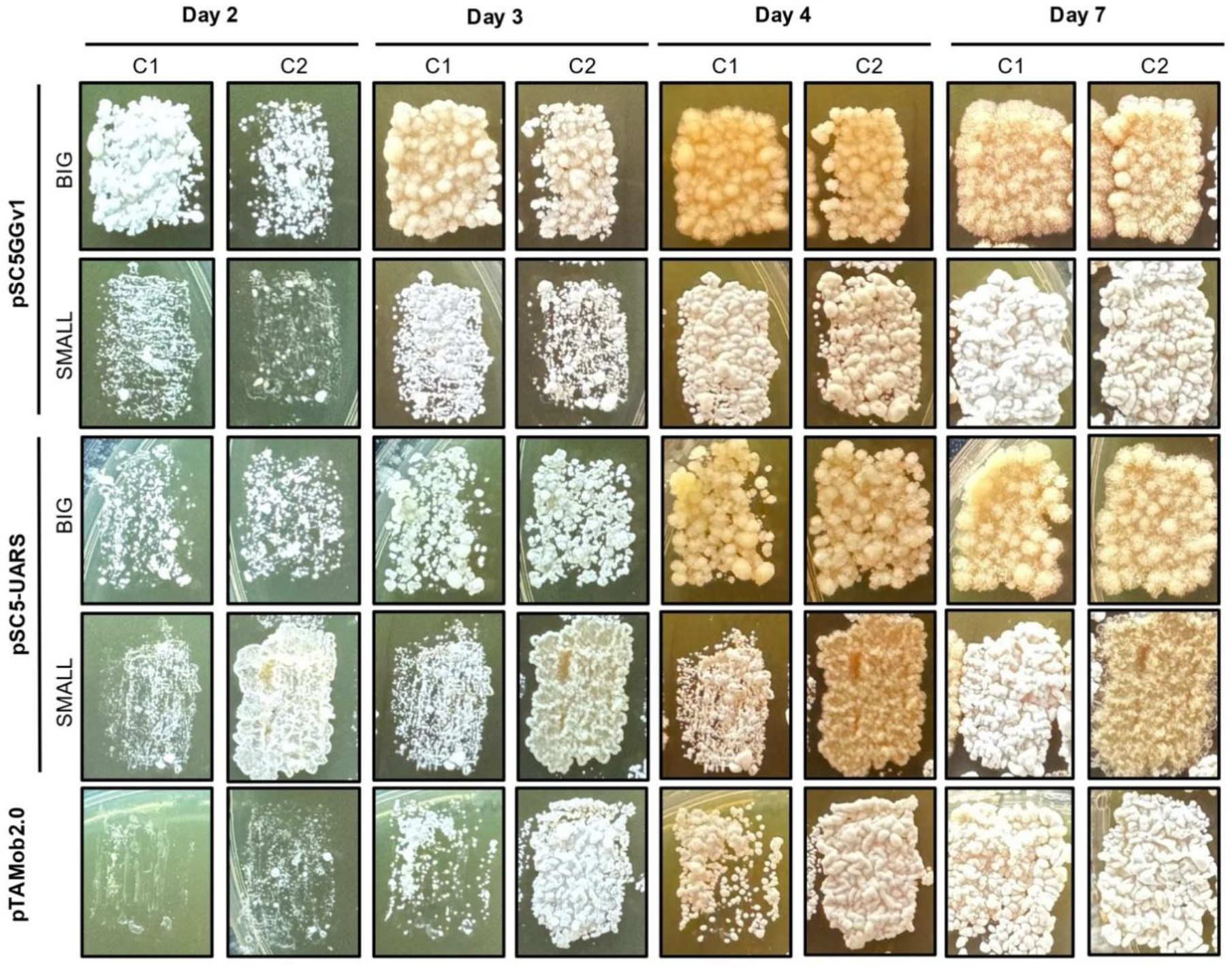
Colonial morphology of *U. maydis* transconjugants re-streaked three times. Two colonies from each of the ‘small’ and ‘big’ morphologies at each of the three plasmid conjugations were struck out on a 2x YPAD NTC 50 µg/mL/AMP 100 µg/mL plate. Timeline of growth displayed, where the plates were imaged at 2, 3, 4, and 7 days.

To validate DNA delivery to *U. maydis* in the transconjugants, the large and small colonies were further evaluated. After re-streaking both sizes of colonies for each plasmid from the NTC50 plates, the clear difference in morphology is maintained (Figure 3). The small colonies from the pSC5GGv1 and pSC5-UARS plates take longer to grow and remain as hard, smooth, and white appearing the most similar to the breakthrough colonies from the pTAMob2.0 plate. In comparison, the big colonies are faster to grow and appear larger, with a fuzzy and yellow colour.

To validate the DNA delivery, genomic DNA isolation was performed for all colonies displayed in Figure 3. Next, a single-plex PCR (Figure 4A) and Multiplex (MPX) PCR screening (Figure 4B) were performed. All ‘big’ colonies from the pSC5GGv1 and pSC5-UARS transconjugants amplified all three single-plex fragments (Figure 4C). The pSC5GGv1 ‘big’ colonies had the expected MPX banding pattern, however the pSC5-UARS ‘big’ transconjugants only demonstrated two out of the three fragments (Figure 4D). Interestingly, in all cases the ‘small’ colonies as well as pTAMob2.0 colonies were negative for either screen. This suggests that the transconjugants can be visually screened based on size for successful DNA delivery.

**Figure 4.**
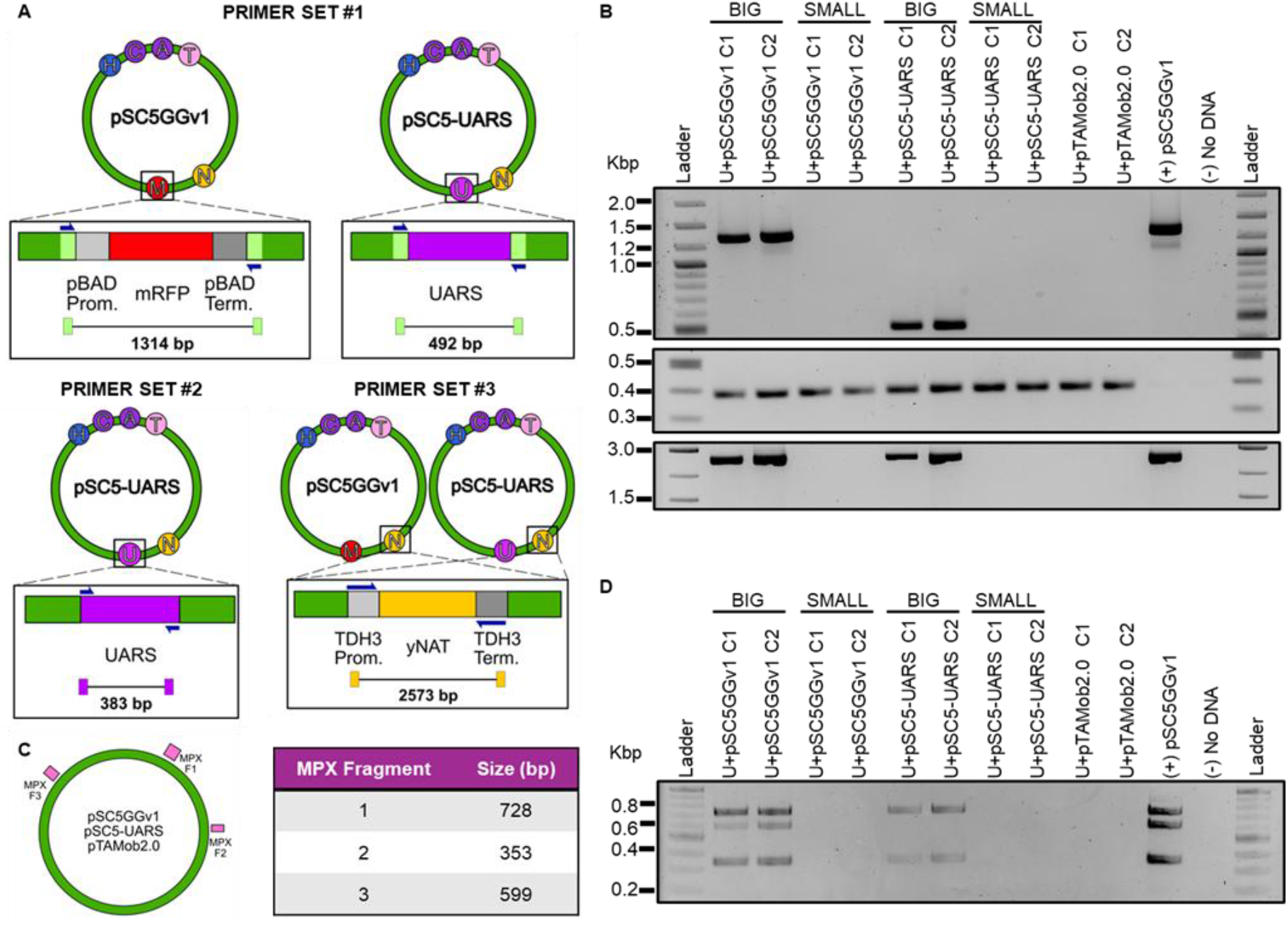
Analysis of *U. maydis* transconjugant colonies. (A) Three single-plex primer sets were designed to confirm DNA delivery. Primer set #1 binds to the same region in pSC5GGv1, and pSC5-UARS highlighted in light green and amplifies different sizes for both plasmids. Primer set #2 amplifies the UARS sequence which only exists in the pSC5-UARS plasmid but is also ubiquitous within the *Ustilago* genome. Primer set #3 amplifies the nourseothricin resistance cassette which only exists in the pSC5GGv1 and pSC5-UARS plasmids. (B) Multiplex primers designed to amplify three regions within all three of the conjugative plasmids. Fragment sizes displayed in the table to the right. (C) Three gels displayed in order of primer sets, and amplifying from each of the two colonies re-streaked for the ‘big’ and ‘small’ morphologies, at each plasmid conjugation. (D) Agarose gel displaying the multiplex screening. All gels were prepared as 2% agarose and ran at 120V for 60 minutes.

We attempted plasmid recovery by transforming the isolated transconjugant DNA into *Epi300 E. coli*. However, in all attempts using both electroporation and chemical transformation we were unable to successfully recover plasmids indicating the integration of the plasmid into the *U. maydis* genome.

## DISCUSSION

DNA delivery to filamentous fungi remains challenging due to their complex cell wall structure and interspecies variability, which necessitate strain-specific optimization (Li et al., 2017). Conjugation, though underexplored, offers advantages including simple preparation and the capacity to transfer large plasmids both *in cis* (all genetic elements required for plasmid transfer and cargo are present on one plasmid) and *in trans* (genetic elements required for plasmid transfer are on separate plasmid form the cargo plasmid). Existing methods such as electroporation, PEG-mediated transformation, and *Agrobacterium*-mediated transformation (AMT) require specialized equipment, extensive optimization, and are limited by DNA size. Here, we report the first demonstration of bacterial conjugation–mediated DNA transfer to the filamentous fungus *U. maydis*.

We evaluated conjugative DNA transfer using the broad-host-range plasmid pSC5GGv1, which has previously been demonstrated to mobilize into several yeast species (Cochrane et al., 2022), along with a variant carrying a *U. maydis* autonomously replicating sequence (UARS) described by (Tsukuda et al., 1988). Using these plasmids, we established an optimized conjugation protocol and confirmed successful DNA transfer to *U. maydis*. Lower antibiotic selection increased colony numbers, though some were breakthrough colonies; but true transconjugants were visually distinguishable. The frequency of transfer into *U. maydis* was lower than that observed for *S. cerevisiae*, but comparable to the levels reported for other non-conventional yeasts. Surprisingly, the inclusion of the UARS element did not enhance colony formation, suggesting that it may be unable to sustain replication of large episomal constructs (56 - 57 kb) or that such plasmids are inherently unstable in *U. maydis*. Additionally, certain sequences within pSC5GGv1 could contribute to toxicity or instability in this host. Future work will explore plasmid size constraints, test conjugation under both cis and trans configurations, and aim to identify additional replication and segregation elements, such as a compatible origin of replication and centromeric sequence, to achieve stable extrachromosomal maintenance.

In summary, this study establishes the first conjugation-based DNA delivery system for *U. maydis*. Further development should focus on constructing conjugative plasmids capable of autonomous replication and enhancing transfer efficiency, potentially through accelerated laboratory evolution (Neil et al., 2021). Optimized conjugative systems could enable programmable DNA delivery, including CRISPR/Cas9-based applications, and provide a framework for antifungal and biocontainment strategies (Jaafar and Carvalhais, 2025).

## AUTHOR CONTRIBUTIONS

TJ: formal analysis, investigation, methodology, writing – original draft, writing – review & editing; AJ: formal analysis, investigation, methodology, writing – review & editing; VN: formal analysis, investigation; methodology, writing – review & editing; BJK: conceptualization, formal analysis, funding acquisition, methodology, resources, supervision, writing – review & editing.

## DATA AVAILABILITY

Data generated or analyzed during this study are available in the published article and its supplementary materials.

