## Supplementary file for "Conjugation-mediated DNA delivery to the filamentous fungus *Ustilago maydis*"

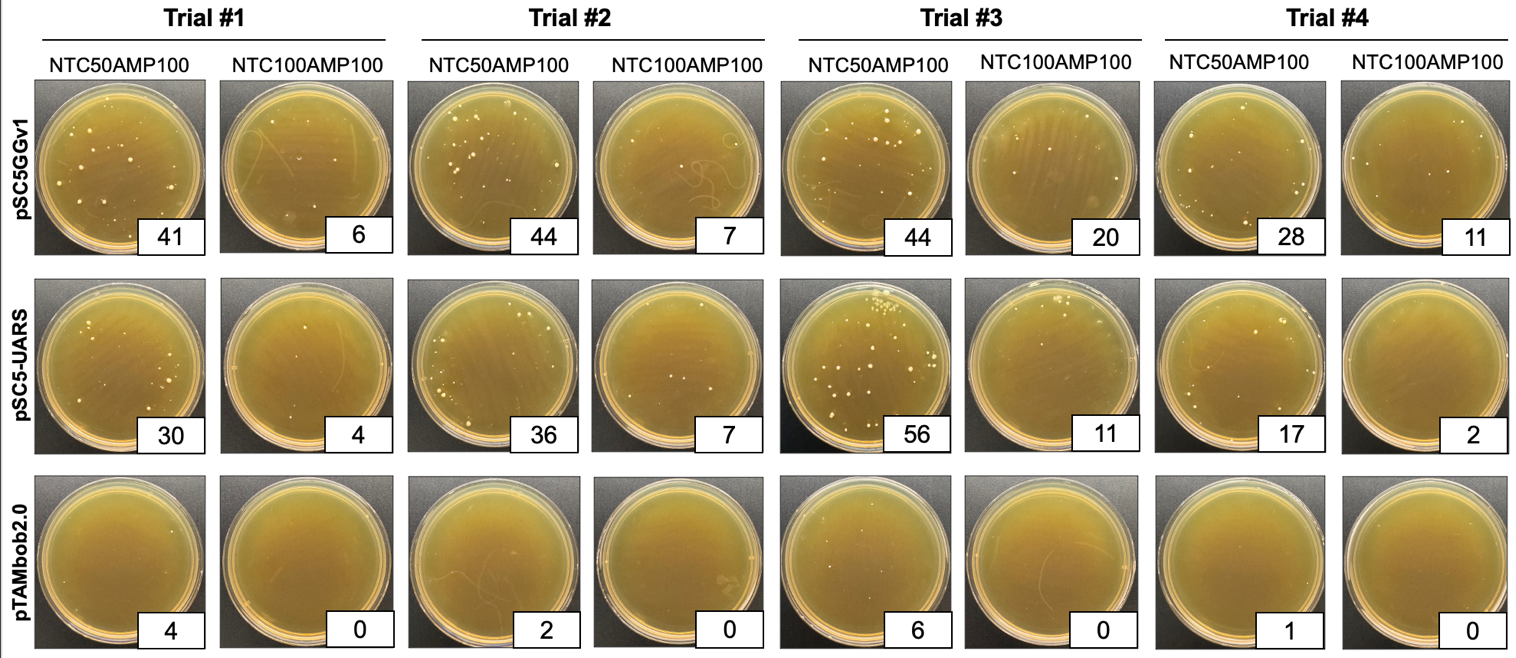

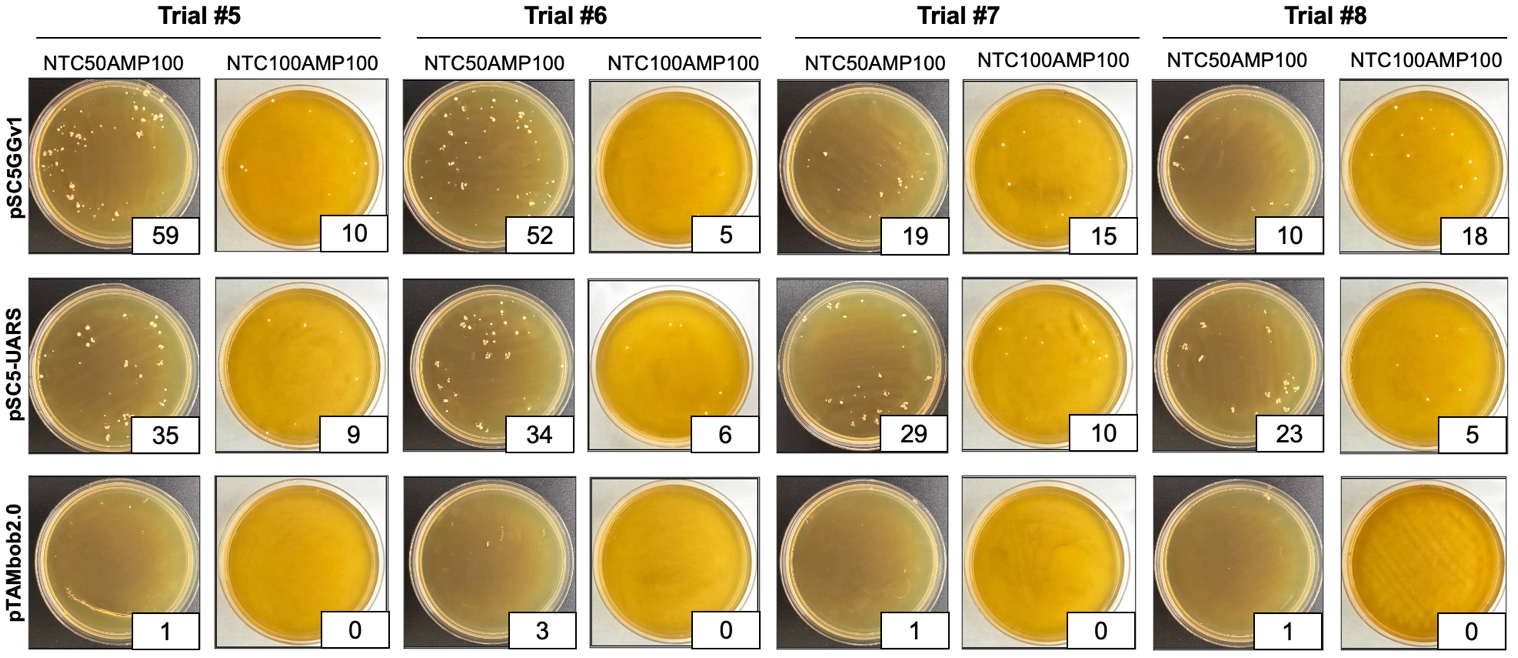

**Figure S1.** Eight biological replicates displaying conjugation of pSC5GGv1, pSC5-UARS, and pTAMob2.0 to *U. maydis* in two different selection plates, NTC50AMP100 and NTC100AMP100. Images taken after 6 days of incubation. Trials 5-8 were imaged with different backgrounds for each of the NTC50, and NTC100 concentrations, causing the difference in plate colouration.

**Table S1.** Primers used to amplify gene inserts from pGL0-21 to create pSC5-UARS. Highlighted in yellow is the part that binds the UARS.

| **FWD Primer** | **REV primer** | **Amplicon Size** |
| --- | --- | --- |
| GGTCTCGGTAAATTGTCATTCGTGATTCTAGTACTGTGGG | GGTCTCGAGCAATTACCGATCCTCGATCTTTGTGCAAGCT | 397 bp |

**Table S2.** Genomic DNA concentration and quality of the two ‘big’ and two ‘small’ colonies isolated from the *U. maydis* transconjugants. Measured using nanodrop spectrophotometer.

|  | **DNA concentration (ng/uL)** | **260/230** | **260/280** |
| --- | --- | --- | --- |
| **U(1) pSC5GGv1-Big C1** | 994.846 | 0.59 | 1.69 |
| **U(1) pSC5GGv1-Big C2** | 417.392 | 0.52 | 1.78 |
| **U(1) pSC5GGv1-Small C1** | 5244.424 | 1.95 | 2.11 |
| **U(1) pSC5GGv1-Small C2** | 3798.241 | 1.93 | 2.07 |
| **U(1) pSC5-UARS-Big C1** | 1259.861 | 0.70 | 1.82 |
| **U(1) pSC5-UARS -Big C2** | 697.122 | 0.89 | 1.73 |
| **U(1) pSC5-UARS -Small C1** | 1039.285 | 1.87 | 1.90 |
| **U(1) pSC5-UARS -Small C2** | 844.481 | 1.21 | 1.90 |
| **U (1) pTAMob2.0 C1** | 3958.908 | 1.82 | 1.99 |
| **U (1) pTAMob2.0 C2** | 3079.320 | 1.42 | 1.98 |

**Table S3.** Primers used in Singleplex screening.

| **Primer Set** | **FWD Primer** | **REV primer** | **Amplicon Size** |
| --- | --- | --- | --- |
| #1 | TCACCACCAGTATAGGCACG | CCGCGCGAGAACCTACTAA | pSC5GGv1 = 1314 bp  pSC5-UARS = 492 bp |
| #2 | ATTACCGATCCTCGATCTTTGTGCAAGCT | ATTGTCATTCGTGATTCTAGTACTGTGGG | 383 bp |
| #3 | TGCTCTGCGAGGCTGGCCGGGTCATGGTTAGATACGGAACC | TAGTTACCGGATAAGGCGCACAAACACCCTTTCAATGGGC | 2613 bp |

**Table S4.** Primers used in Multiplex screening.

| **MPX Fragment** | **FWD Primer** | **REV primer** | **Amplicon Size** |
| --- | --- | --- | --- |
| #1 | CACTGAAGACTGCGGGATTG | TGAGAGCAGGAAGAGCAAGA | 728 bp |
| #2 | GCCGAATATTTACGCCAGGG | GTTGCGGGCTTCTTCTTGAA | 353 bp |
| #3 | CCGTGATTGATGATATAGCGGC | CGCCAAGCTGCAATTCCTT | 2613 bp |

**Table S5. Anova summary**

|  | **F-value** | **Num DF** | **Den DF** | **Pr > F** |
| --- | --- | --- | --- | --- |
| **Plasmid** | 46.1665 | 2.0000 | 14.0000 | 0.0000 |
| **Concentration** | 28.6159 | 1.0000 | 7.0000 | 0.0011 |
| **Plasmid: Conc.** | 8.9854 | 2.0000 | 14.0000 | 0.0031 |

**Table S6.** Paired t-tests summary.

| **Comparison** | **t** | **p-value** | **Bonferroni corrected p-value (multiplied by 9)** |
| --- | --- | --- | --- |
| **pSC5GGv1 vs pSC5-UARS (NTC50)** | 0.9095785621190150 | 0.39328494174074400 | 1.0 |
| **pSC5GGv1 vs pTAMob2.0 (NTC50)** | 6.1031029486452700 | 0.0004895824649314990 | 0.004406242184383490 |
| **pSC5-UARS vs pTAMob2.0 (NTC50)** | 8.443341921006170 | 6.44912786453402E-05 | 0.0005804215078080610 |
| **pSC5GGv1 vs pSC5-UARS (NTC100)** | 2.4737374205250500 | 0.042599092523041000 | 0.3833918327073690 |
| **pSC5GGv1 vs pTAMob2.0 (NTC100)** | 4.28757721629013 | 0.003621462218259620 | 0.03259315996433660 |
| **pSC5-UARS vs pTAMob2.0 (NTC100)** | 6.14817045957576 | 0.00046837779076746300 | 0.004215400116907170 |
| **pSC5GGv1 (NTC50 vs NTC100)** | 3.0198981202677600 | 0.019389104591877800 | 0.17450194132690000 |
| **pSC5-UARS (NTC50 vs NTC100)** | 7.818043917732140 | 0.00010554171791328600 | 0.0009498754612195780 |
| **pTAMob2.0 (NTC50 vs NTC100)** | 3.63735707515039 | 0.008316210450525760 | 0.07484589405473180 |

**Table S7.** Colony counts of conjugation to *U. maydis* using different ratios of fungi to bacteria. The numbers displayed in the ratio are the volume in uL.

| Plasmid | U100:B200 | U100:B100 | U200:B100 |
| --- | --- | --- | --- |
| pSC5GGv1 | 14 | 3 | 7 |
| pSC5-UARS | 4 | 3 | 1 |
| pTAMob2.0 | 0 | 0 | 0 |

**Table S8. Colony Counts on rich vs. poor media**

| Plasmid | Rich Media (2x YPAD) | Poor Media (-HIS) |
| --- | --- | --- |
| pSC5GGv1 | 10 | 3 |
| pSC5-UARS | 36 | 5 |
| pTAMob2.0 | 0 | 0 |
